# Power spectrum slope confounds estimation of instantaneous oscillatory frequency

**DOI:** 10.1101/2021.06.21.449312

**Authors:** Jason Samaha, Michael X Cohen

**Affiliations:** Psychology Department, University of California, Santa Cruz; Donders Centre for Medical Neuroscience, Radboud University Medical Centre

**Keywords:** instantaneous frequency, neural oscillations, alpha-band, 1/f, simulations

## Abstract

Oscillatory neural dynamics are highly non-stationary and require methods capable of quantifying time-resolved changes in rhythmic activity in order to understand neural function. Recently, a method termed ‘frequency sliding’ was introduced to estimate the instantaneous frequency of oscillatory activity, providing a means of tracking temporal changes in the dominant frequency within a sub-band of field potential recordings. Here, the ability of frequency sliding to recover ground-truth oscillatory frequency in simulated data is tested while the exponent (slope) of the 1/*f*^x^ component of the signal power spectrum is systematically varied, mimicking real electrophysiological data. The results show that 1) in the presence of 1/*f* activity, frequency sliding systematically underestimates the true frequency of the signal, 2) the magnitude of underestimation is correlated with the steepness of the slope, suggesting that, if unaccounted for, slope changes could be misinterpreted as frequency changes, 3) the impact of slope on frequency estimates interacts with oscillation amplitude, indicating that changes in oscillation amplitude alone may also influence instantaneous frequency estimates in the presence of strong 1/*f* activity; and 4) analysis parameters such as filter bandwidth and location also mediate the influence of slope on estimated frequency, indicating that these settings should be considered when interpreting estimates obtained via frequency sliding. The origin of these biases resides in the output of the filtering step of frequency sliding, whose energy is biased towards lower frequencies precisely because of the 1/*f* structure of the data. We discuss several strategies to mitigate these biases and provide a proof-of-principle for a 1/f normalization strategy.

## Introduction

Decades of electrophysiological research has unveiled systematic relationships between properties of rhythmic brain activity and behavior, spurring myriad theoretical accounts of the role that neural oscillations play in perceptual, cognitive, and motor processes (Bastos et al., 2012; A. Engel & Fries, 2010; A. K. Engel et al., 2001; Foxe & Snyder, 2011; Fries, 2005; Jensen & Mazaheri, 2010; Jones, 2002; Klimesch et al., 2007; Mathewson et al., 2011; Palva & Palva, 2011; Samaha et al., 2020; Spitzer & Haegens, 2017; VanRullen, 2016; Varela et al., 1981). By and large, the focus of most work has been on amplitude and phase dynamics of brain rhythms. However, several findings have highlighted the importance of peak oscillation frequency within a particular band for shaping behavior and neural processing (Angelakis et al., 2004; Atallah & Scanziani, 2009; Cohen, 2014; Dipoppa & Gutkin, 2013; Furman et al., 2018; Haegens et al., 2014; Klimesch et al., 1993; Mierau et al., 2017; Nelli et al., 2017). For instance, within- and between-subject variation in alpha-band frequency (7-14 Hz) is predictive of temporal properties of visual (Baumgarten et al., 2018; S. Coffin & Ganz, 1977; Stephen Coffin, 1977; Gray & Emmanouil, 2020; Gulbinaite, İlhan, et al., 2017; Kristofferson, 1967; May et al., 2014; Minami & Amano, 2017; Ro, 2019; Samaha & Postle, 2015; Shen et al., 2019) as well as cross-modal perception (Cecere et al., 2015; Cooke et al., 2019; Keil & Senkowski, 2017)

Often, peak frequency is computed as the single frequency with largest amplitude within a sub-band of the power spectrum (Angelakis et al., 2004; Cecere et al., 2015; Grandy et al., 2013; Gulbinaite, Viegen, et al., 2017; Samaha & Postle, 2015), or as the ‘center of mass’ of the whole oscillation peak in the power spectrum (Furman et al., 2018; Jann et al., 2010; Klimesch et al., 1993), or by fitting a Gaussian function to peaks in the power spectrum (Donoghue et al., 2020; Haegens et al., 2014; Jin et al., 2006). However, these spectrum-based methods assume that the dominant frequency of a sub-band is approximately stationary over time, as the power spectrum represents signal properties collapsed over a given time window. Recently, a method for characterizing instantaneous changes in the dominant frequency of a signal was introduced (Cohen, 2014), building upon mathematical models if instantaneous frequency estimation (Boashash, 1992). Termed *frequency sliding*, time-resolved changes in frequency are estimated by computing the temporal derivative of the phase angle timeseries obtained after Hilbert-transforming a narrowband-filtered signal.

Frequency sliding has been used in a number of experiments to demonstrate temporal fluctuations in oscillatory frequency in response to varying stimulus properties (Cohen, 2014; Gulbinaite et al., 2019; Noguchi & Kubo, 2020), task demands (Wutz et al., 2018), as a predictor of trial-to-trial variability in perception (Nelli et al., 2017; Samaha & Postle, 2015; Shen et al., 2019), and as a method for computing functional connectivity between brain areas (Cohen, 2014). Despite adoption in the field, the method has not been extensively tested in the recovery of ground-truth oscillatory frequency. Initial simulations were presented in the paper that introduced the method in the context of electrophysiology data (see Figure 1B from Cohen, (2014)), which revealed that the use of a band-pass filter with a plateau shape in the frequency domain (the first step of the frequency sliding analysis; see methods) did not bias the estimated frequency towards the center of the filter band (as, presumably, a filter with a Gaussian response in the frequency domain would). However, frequency sliding has not been evaluated in the context of changing physiologically-relevant parameters of the power spectrum. In particular, it has recently been noted how the exponent (i.e., slope) of the 1/*f*^x^ component of the signal power spectrum (also referred to as the *aperiodic* component) can, if not accounted for, confound measurements such as oscillatory power or band-power ratios (Donoghue et al., 2020, 2020). Moreover, aperiodic slope has been found to vary considerably both within- (Donoghue et al., 2020; Podvalny et al., 2015) and between-individuals (Donoghue et al., 2020; Voytek et al., 2015), particularly across development (Schaworonkow & Voytek, 2020). Because oscillatory activity is embedded within such aperiodic activity, an investigation of whether aperiodic exponent variation impacts estimation via frequency sliding is warranted.

**Figure 1.**
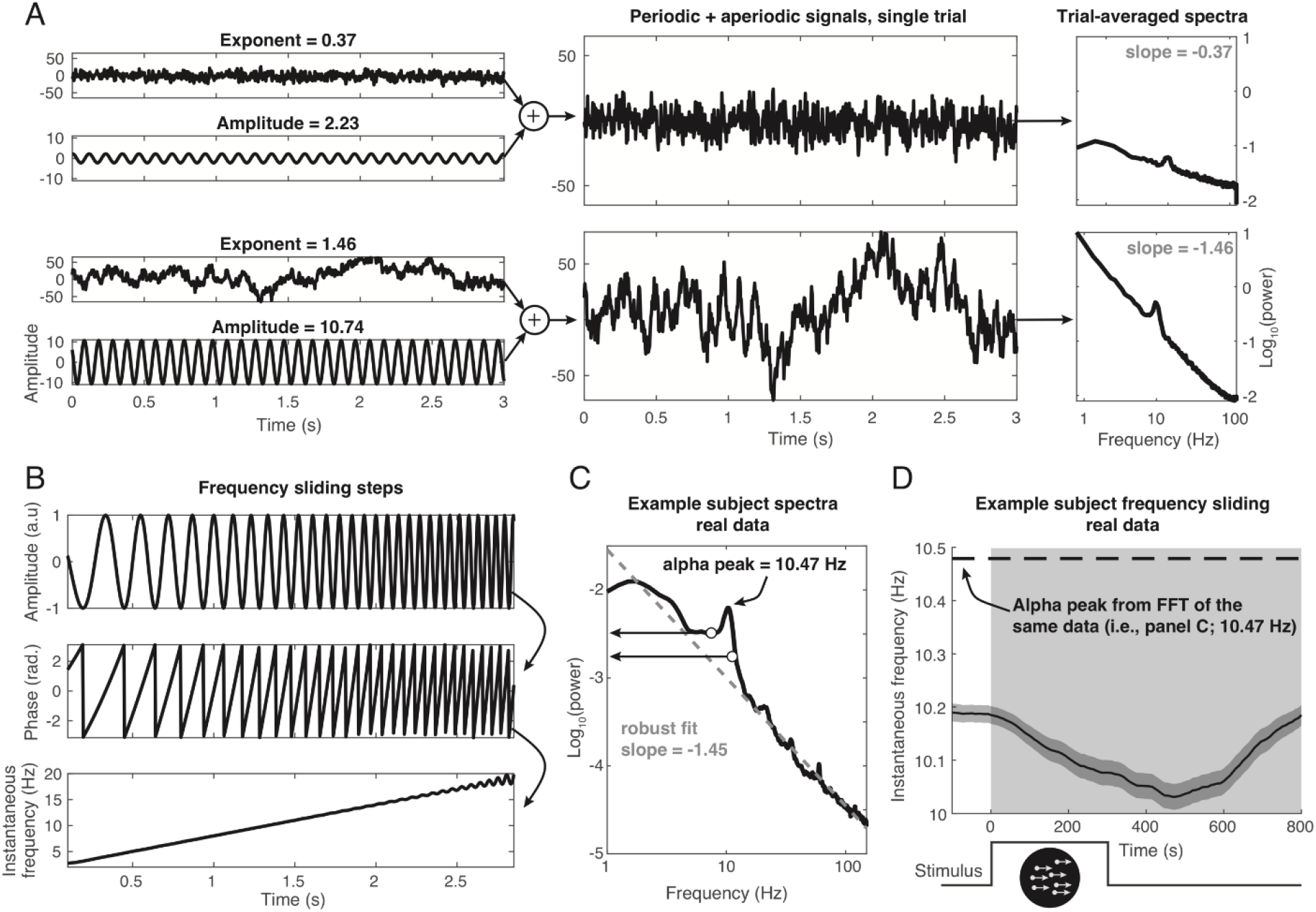
Simulation/recovery method and the underestimation of instantaneous oscillation frequency in real data. **A**) Simulated signals contained an aperiodic 1/*f*^x^ component with a variable exponent and a periodic (sine wave) component with a fixed frequency of 10 Hz but a variable amplitude. The top panel shows an example trial with a relatively low exponent (shallow slope) and low amplitude. The bottom panel shows an example trial with relatively high exponent (steep slope) and medium-high amplitude. Signals were scaled to approximate the microvolt range observed in single-trial EEG recordings and summed. Example log-log spectra on the right are averaged over 10,000 simulated trials and show the effect of varying parameters in the timeseries. **B**) Frequency sliding (Cohen, 2014) can be used to recover instantaneous fluctuations in oscillatory frequency. Here, the logic is shown for a cosine signal increasing linearly in frequency from 2 to 20 Hz over the course of 3 seconds (a “chirp”). Once the band of interest in the signal has been isolated (typically via filtering), the phase angle is extracted via Hilbert transform (second panel), and its temporal derivative is computed and scaled to Hz (last panel). In this example, frequency sliding accurately captures the temporal changes in frequency. **C**) Peak frequency estimated via FFT of real EEG data recoded from the first author’s scalp (electrode POz) averaged over trials of a dot-motion direction discrimination task (n=973; 0-800 ms post-stimulus). The log-log spectrum shows a strong negative slope (grey dashed line; exponent approx. ~1.45, cf. spectrum in panel A) obtained via robust linear fit. The peak alpha frequency is at 10.47 Hz. Notice how, because of the 1/f slope, the lower edge of the alpha band (8 Hz) has greater power than the upper edge of the band (12 Hz), indicated with circles and arrows. **D**) Frequency sliding results of these data show a clear underestimation of the peak frequency obtained via FFT (black dashed line). The gray window spans timepoints used for the FFT in panel C; shaded bands denote ± 1 SEM across trials; inset below denotes timing of the dot-motion stimulus.

To preview, the attempt to recover simulated oscillatory frequency revealed a systematic underestimation of the true oscillation frequency, the magnitude of which depended on the slope the power spectrum and the amplitude of the oscillation. Thus, keeping oscillatory frequency constant and varying aperiodic slope or oscillation amplitude could manifest in changes to the instantaneous frequency estimated via frequency sliding, producing a possible confound in real data. Although frequency sliding is a valuable tool for uncovering time-varying frequency modulations in electrophysiological data, users of the method should be aware of these alternative causes of estimated frequency modulation in their data, and future work should examine ways of attenuating the confounds uncovered in the present simulations. We discuss several possible mitigating strategies.

## Methods

### Simulated data

Signals were simulated in the MATLAB environment (2020a, version 9.8.0) with use of the MATLAB DSP System Toolbox. The code and simulated signals are freely provided at https://osf.io/f8jqd/. Simulated signals comprised a periodic (sinewave) and an aperiodic (1/*f*) component, which were first independently generated and then summed. Details of each signal component are described below.

The aperiodic component of the power spectrum was simulated using the *dsp. ColoredNoise* MATLAB object, which generates random values that follow a 1/*f*^x^ power spectrum, with the exponent, *x*, as a controllable parameter. Signals were simulated with 8 different exponents ranging linearly from 0.1 to 2 (flatter to steeply negative). Note that the exponent corresponds to the slope of a straight line fit to the aperiodic power spectrum in log-log coordinates multiplied by −1. The range of exponents simulated here was chosen to encompass the distribution of actual exponents measured during resting-state in a large sample of adults (Donoghue et al., 2020), which has a mean of 0.83, but which can vary considerably across the first months of development (Schaworonkow & Voytek, 2020). Lastly, the aperiodic signal was mean-centered and scaled by a factor of 9 to bring it within range of typical single-trial EEG data (approx. −40 to 40 microvolts when *x =* 0.91). Example single-trial signals are shown in Figure 1A. With these parameters, the “rotation point” in the power spectrum – that is, the frequency about which the spectrum pivots – is around 45 Hz (see Figure 2). In intracranial recordings, the rotation point was found to vary widely across brain areas, but has a modal value around 30 Hz in visual areas, 40 Hz in auditory areas, and 60 Hz in motor areas (Podvalny et al., 2015). Thus, the aperiodic activity simulated here has many features in common with empirical electrophysiological spectra.

**Figure 2.**
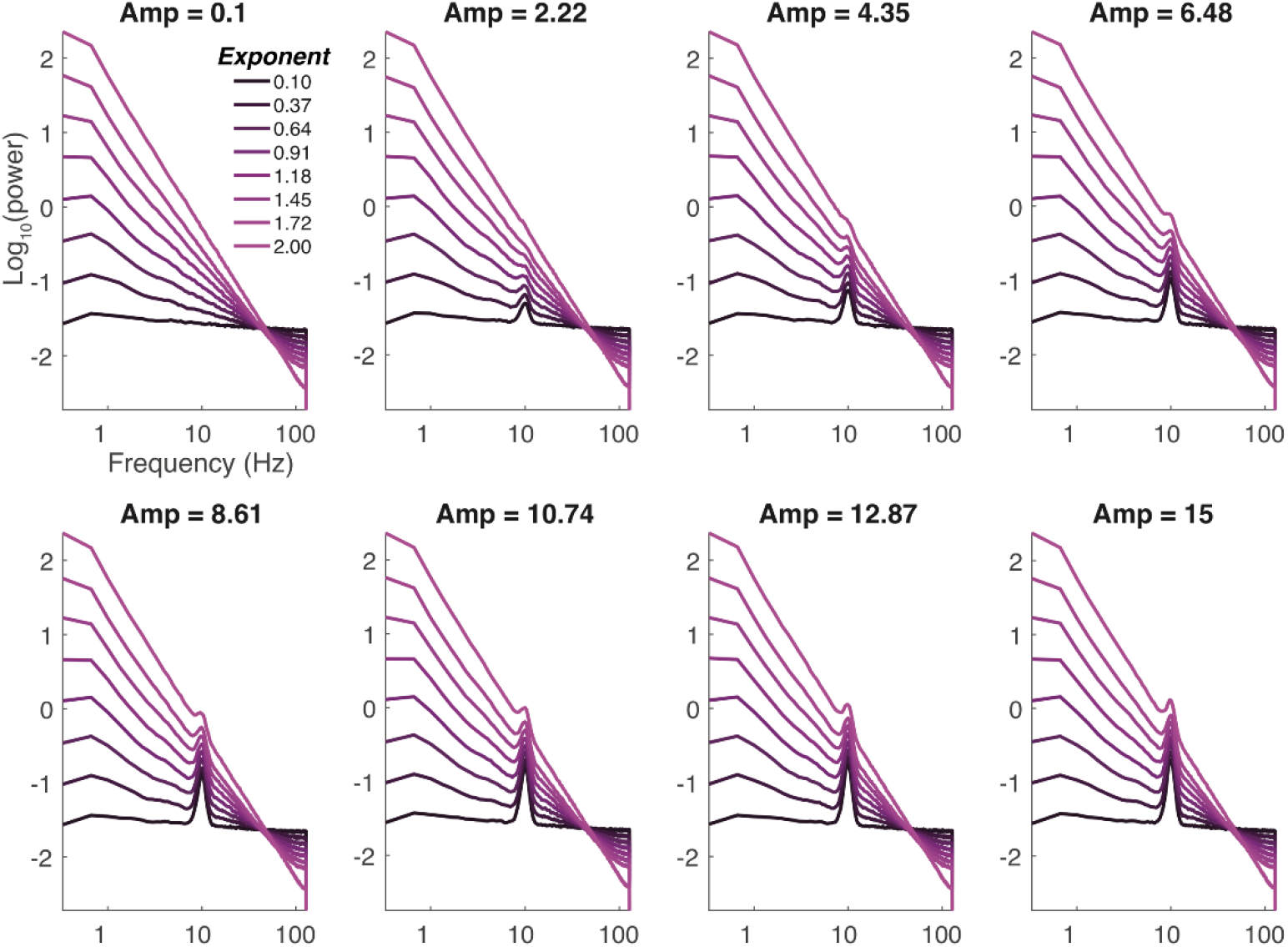
Power spectra of the simulated signals averaged over trials (trial duration = 3 seconds; n=10,000 in each spectrum). Each spectrum shows a unique combination of periodic amplitude and aperiodic exponent (−1*slope). Coordinates are log-log spacing.

Sine waves (i.e., oscillations or the periodic component) were generated and summed with the aperiodic signal. Sine waves were defined with a mean frequency of 10 Hz and trial-to-trial standard deviation (SD) of 1 Hz in order to approximate the typical bandwidth of alpha activity in real data (Donoghue et al., 2020; Haegens et al., 2014). The choice to model the human alpha rhythm is motivated by the fact that frequency sliding has been applied particularly often to the study of alpha frequency (Cohen, 2014; Gulbinaite, Viegen, et al., 2017; Nelli et al., 2017; Samaha & Postle, 2015; Shen et al., 2019; Wutz et al., 2018). Oscillation amplitude varied across 8 levels, linearly spaced between 0.1 to 15 (small to large). This range approximates the range of peak-to-peak amplitudes of single-trial alpha rhythms observed in EEG, measured in microvolts. The phase of the sinewave varied randomly from trial to trial between -pi and pi, to mimic spontaneous (i.e., non-phase-locked) activity.

In total 10,000 trials were simulated at each of the 8 exponent levels and 8 amplitude levels, for a total of 640,000 simulated trials. Each trial was 3 seconds long, with a sampling rate of 256 Hz. To avoid filter-related edge artifacts in plots and averages, only the central second of each trial (i.e., 1-2 seconds) was extracted following the frequency sliding analysis. Power spectra were constructed by applying a fast Fourier transform (*fft.m*) to the detrended, Hamming-tapered, data from each 3 second epoch. The absolute value of the resulting Fourier coefficients were squared and log_10_ transformed (on single trials; see (Smulders et al., 2018) to obtain power in the frequency domain.

### Frequency sliding

Instantaneous frequency was estimated following the frequency sliding method described in Cohen, (2014). Briefly, the simulated signals (comprising the sum of the periodic and aperiodic component) were narrowband filtered between 8 and 12 Hz using a FIR (*firls.m*) filter with a plateau-shaped frequency response in order to prevent biasing the filtered data towards the center of the band. The filter had a transition width of 15% of the lower and upper frequency bounds and was applied in the forward and reverse direction (*filtfilt.m*) to achieve a zero-phase filter. A Hilbert transform (*hilbert.m*) was applied to the narrowband filtered data and the phase angle time series was extracted (*angle.m*). Lastly, the phases were unwrapped (*unwrap.m*), and the first temporal derivative was computed (*diff.m*). When scaled by the sampling rate and 2π, the temporal derivative of the phase angle time series estimates the instantaneous frequency in the filtered band (see Figure 1B).

To avoid phase slips or other sudden transitions in the phase time series from causing large spikes in the instantaneous frequency estimate, the scaled derivative was filtered 10 times with median filters spanning 10 to 400 ms. The median filter step is less critical in the simulated data, but was included to keep the analysis close to the protocol for real data, where non-physiological artifacts need to be attenuated via this median filtering protocol (Cohen, 2014; Gulbinaite, Viegen, et al., 2017; Samaha & Postle, 2015; Wutz et al., 2018).

A filter range of 8 to 12 Hz was chosen as it is centered on the simulated oscillation of 10 Hz, encompasses the full alpha peak (given the SD of the simulated peaks), and is a commonly used definition of the alpha-band. Different filter ranges were explored, but unless otherwise stated, the default was 8 to 12 Hz. In plots, the 95% confidence interval (CI) on outcome measures are presented in order to give a sense of the variability across the simulations rather than to infer any statistical significance given that the CI could be arbitrarily scaled by changing the number of simulated trials.

### Correction for 1/f

Our attempt at removing the frequency sliding bias imposed by the 1/*f* was based on constructing an “anti-1/*f*-modulator.” Although there are several naïve and adaptive methods for removing the 1/*f* component of the power spectrum (Donoghue et al., 2020; Groppe et al., 2013; Hughes et al., 2012; Wen & Liu, 2016) these methods are based on remaining in the frequency domain. This provides greater freedom, for example, by allowing relatively negative power values. However, our goal requires time-domain data with the 1/*f* removed, and thus the demodulated power spectrum must be non-negative.

We fit the power spectrum (in log_10_-log_10_ space) using a polynomial regression. The order was determined via the Bayes Information Criteria on the sum or squared errors from model orders ranging from 1 to 5. The inverse of the best-fit model (that is 1/ŷ) was then multiplied (frequency-wise) by the empirical power spectrum. The empirical power spectrum scaled by the anti-1/*f*-modulator is relatively flat, while still preserving the local features (see Figure 6). Finally, the modulated power spectrum was combined with the original phase spectrum, and the inverse FFT was taken to obtain a time-domain signal.

**Figure 3.**
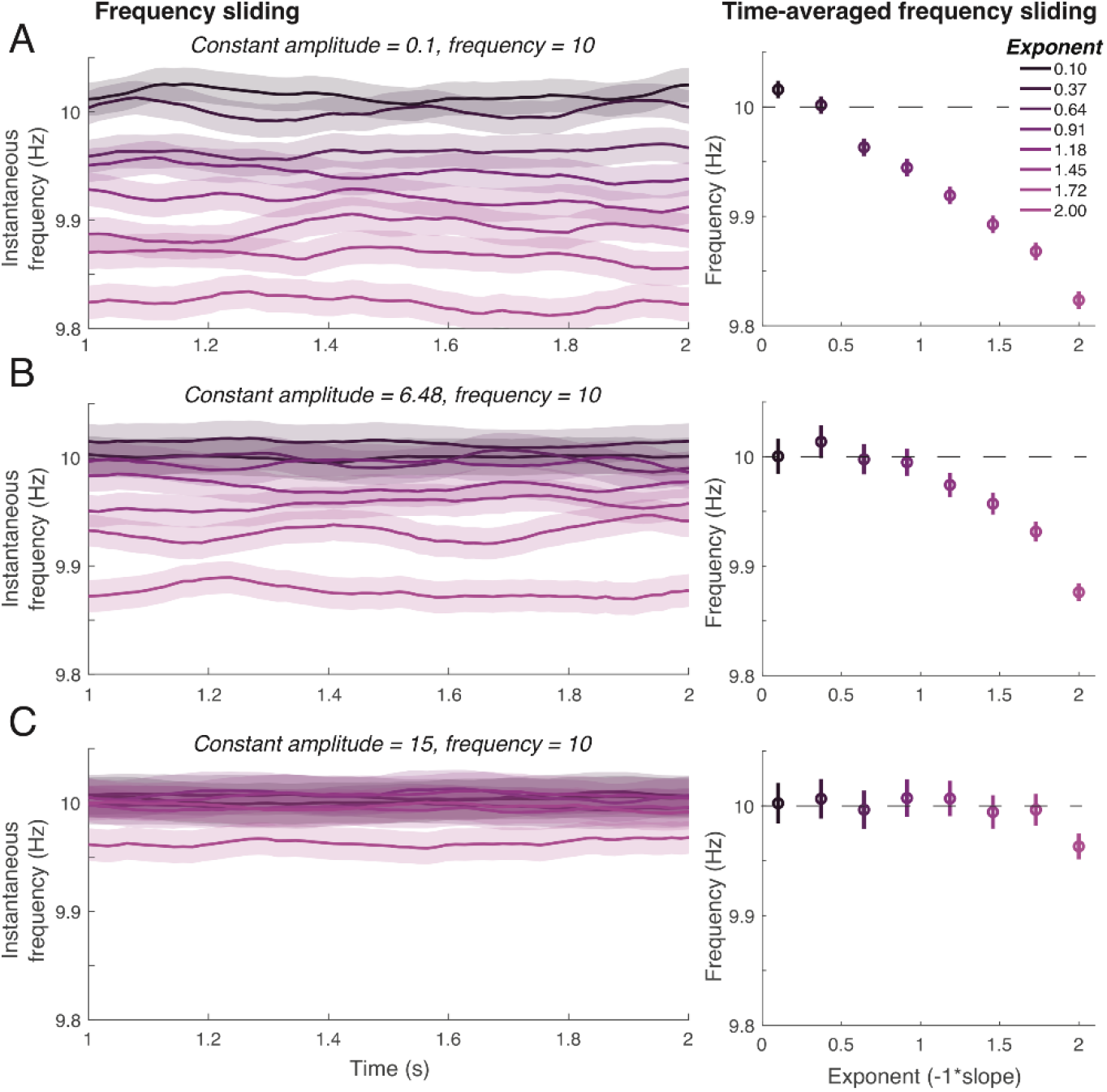
Accuracy of recovering simulated frequency via frequency sliding. The left plot in each panel shows the time course of frequency estimates whereas the right plot shows the same data averaged across time. The effect of varying the aperiodic exponent on estimated frequency is shown for 3 levels of oscillation amplitude: low, medium, and high **A**) At low amplitudes, frequency is increasingly biased as the slope of the aperiodic component of the power spectrum increases. **B**) For a medium amplitude oscillation, underestimation is attenuated, particularly at shallow slopes (small exponents), but still notable at medium-to-large exponents. **C**) At the largest level of amplitude simulated, the impact of the aperiodic slope was nearly absent, discernable only at the steepest slope simulated. Shaded bands and error bars denote the 95% CI across simulations.

**Figure 4.**
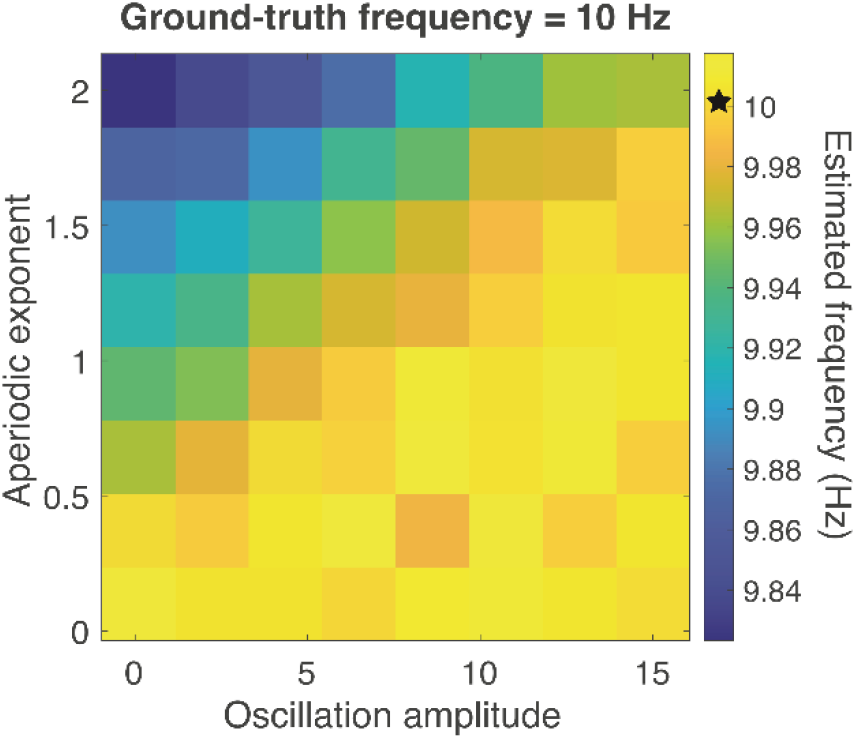
Pseudocolor plot showing the joint effects of varying oscillation amplitude and aperiodic exponent on estimated frequency. Ground truth frequency was 10 Hz (denoted by the star on the color scale). Both parameters influence estimated frequency independently and interactively, with the maximal bias occurring when a power spectrum is steep, and an oscillation is low in amplitude.

**Figure 5.**
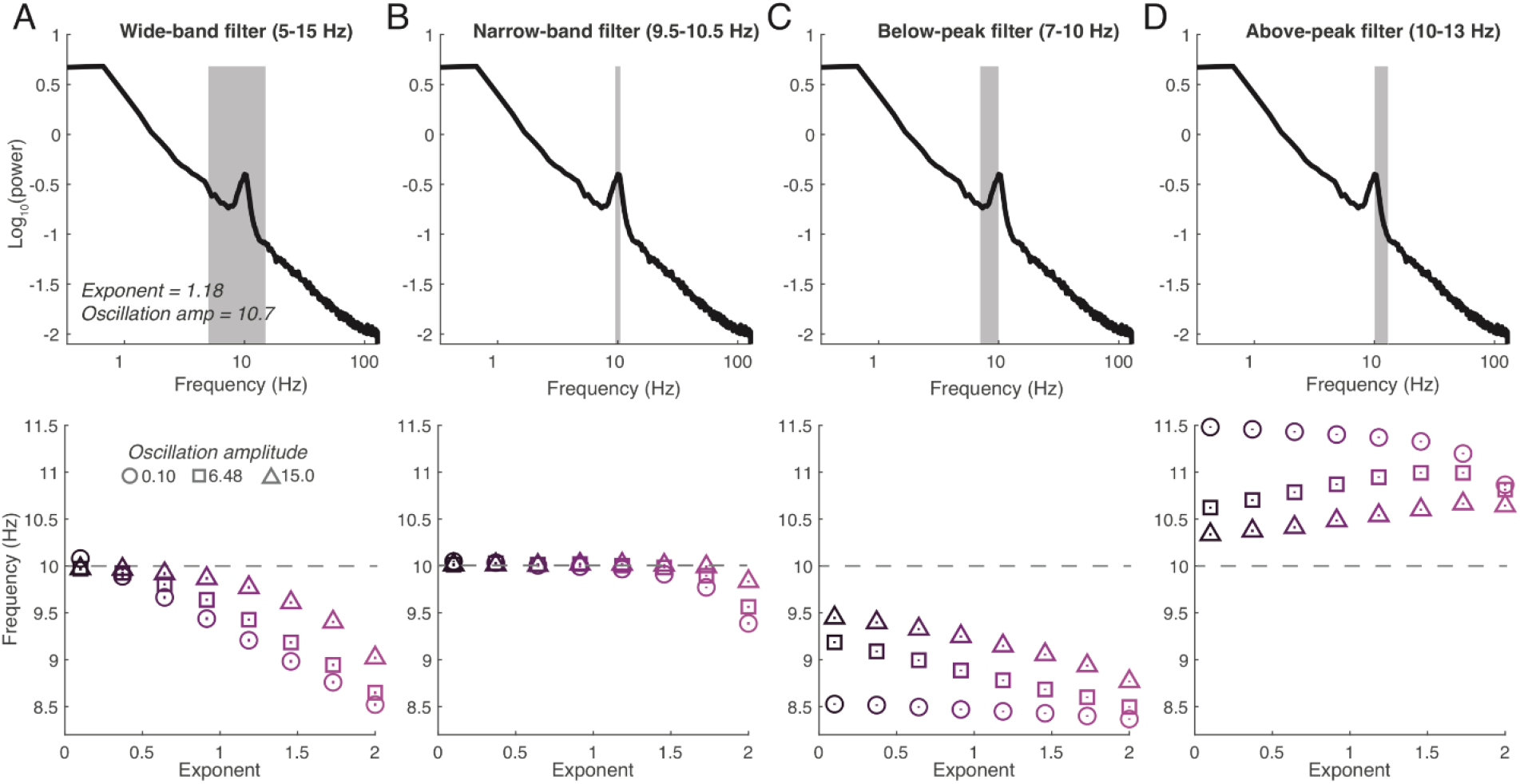
The impact of filter bandwidth and location on the relationship between oscillation amplitude, aperiodic slope, and estimated frequency. Frequency sliding was performed on the same set of simulated data but with varying filter parameters. In each panel, the top plot shows the filter settings on an example spectrum, and the bottom plot shows the estimated frequency at all levels of simulated exponent, and three levels of oscillation amplitude: small, medium, and large. **A**) Example of a wide-band filter, which a researcher may use if they expect frequency to vary widely or if they are unsure of the underlying oscillation frequency and attempt to ‘cast a wide net’. Setting a filter band that is centered on the true oscillation peak (10 Hz) but extends substantially below and above that peak will dramatically increase the influence of the aperiodic component, leading to amplitude-dependent underestimations of up to ~1.5 Hz. **B**) An overly-narrow filter can sufficiently suppress the 1/f influence, but would then presumably be relatively insensitive to any fluctuations in frequency (see (Cohen, 2014)), undermining the purpose of using frequency sliding in the first place. Dashed line in lower plots indicates the ground-truth frequency (10 Hz). A filter placed below (**C**) or above (**D**) the true peak frequency might occur if the researcher uses a default band for all subjects when individual differences in peak frequency exist. A below-peak filter leads to strong, amplitude-dependent underestimation (owing to a bias towards lower frequencies) whereas an above-peak filter leads to amplitude-dependent overestimation. In nearly all cases, misestimation is greater when oscillatory amplitude is low.

**Figure 6.**
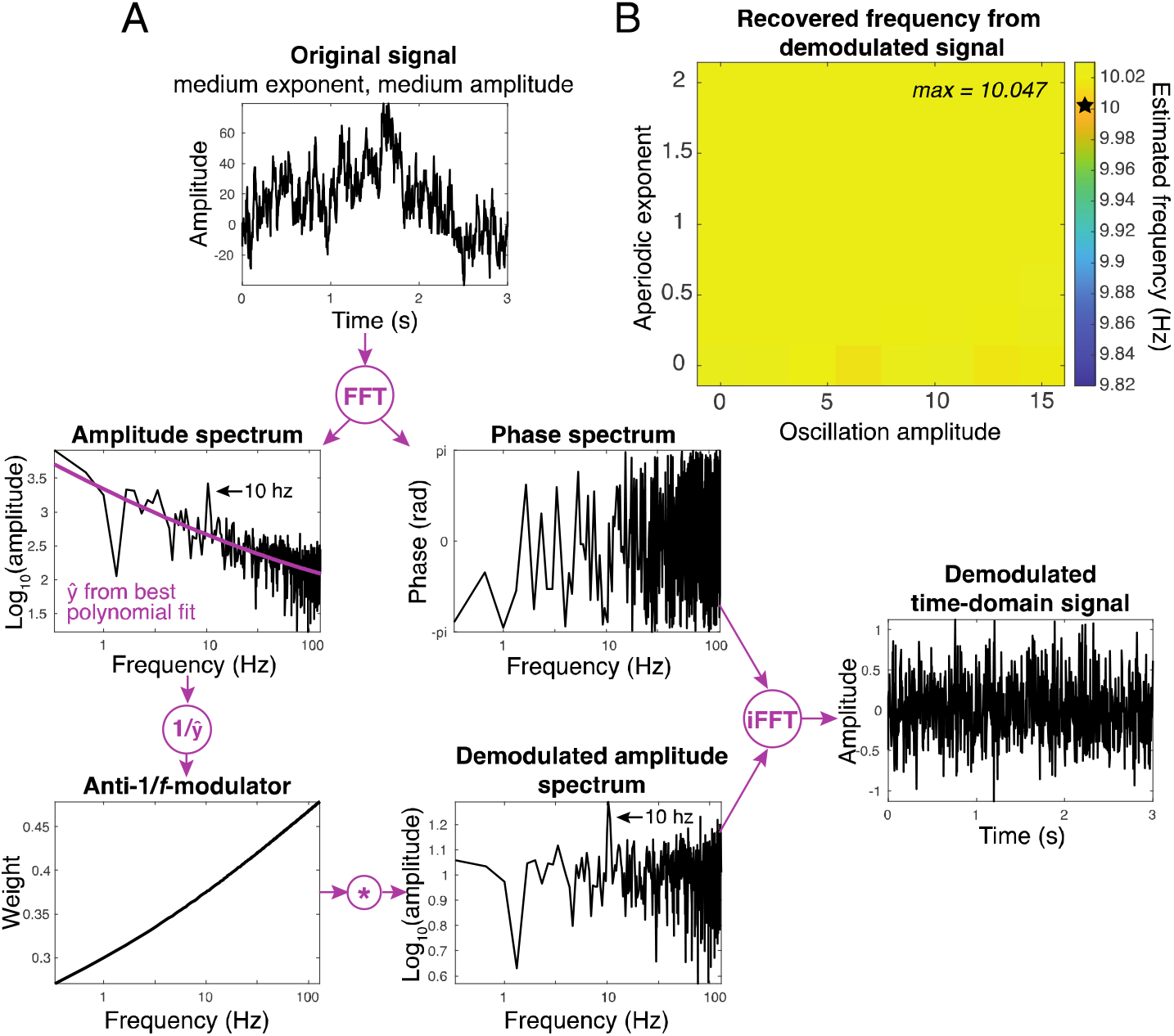
Reconstructing the time-domain signal with attenuated 1/f. **A**) Algorithm for removing aperiodic activity from the time-domain signal in an example simulated trial. Each step in the method is shown in purple arrows and font. First, an FFT is used to obtain the amplitude and phase spectrum of the signal. The amplitude spectrum (in log-log space) was fit with a polynomial, the reciprocal of which was then pointwise multiplied (asterisk) with the amplitude spectrum to obtain a demodulated spectrum. The demodulated spectrum was combined with the original phase spectrum to create complex values that were then input into and inverse FFT to recover a demodulated time-domain signal. Note that the demodulated signal contains much higher energy in high frequencies, but the 10 Hz peak is preserved. **B**) Trial-averaged frequency sliding for each simulated exponent and oscillation amplitude shown on a similar color scale as Figure 4 for visual comparison. Removing the 1/f from the signal resulted in frequency estimates less biased by slope and amplitude, though a slight overestimation was present (max = 10.047 Hz).

## Results

The logic of frequency sliding is demonstrated in Figure 1B as applied to a chirp signal (a cosine wave increasing in frequency from 2 to 20 Hz over the course of 3 seconds). As can be seen, frequency sliding accurately estimates the true frequency of the signal. When applied to data with a 1/*f* structure, however, underestimation can occur. Figure 1C shows an example power spectrum recorded from electrode POz of the first author’s scalp, averaged over 973 trials of visual dot-motion direction discrimination task (FFT from 0 to 800 ms post-stimulus, stimulus duration = 300 ms). The spectrum has an approximate exponent of 1.45, which is within the higher end of the signals simulated here. (For illustration, the slope was estimated via a line fit using the robust bisquare method in log-log units to downweigh the influence of oscillation peaks; for a more robust parameterization method, see (Donoghue et al., 2020)). The peak alpha frequency (frequency with maximum amplitude) during stimulus processing was 10.47 Hz. However, frequency sliding systematically underestimated the FFT-based peak frequency throughout the entire trial period (mean ± SEM across trials of the frequency estimated from frequency sliding during the 800 ms post-stimulus = 10.09 ±0.013).

This underestimation may simply be a product of the FFT-peak detection method being insensitive to the full shape of the peak in spectral energy (as it is based only on a single data point) – indeed, this has been a motivation for other approaches to peak estimation, such as Gaussian fitting (Donoghue et al., 2020; Haegens et al., 2014; Jin et al., 2006) or center-of-mass estimation (Klimesch et al., 1993). However, we are concerned with the underestimation of frequency sliding relative to the FFT-derived peak, which we believe occurred because the lower end of the filter band (here, 8 Hz) has higher spectral energy than the upper end (12 Hz) due to the 1/f background. This is visualized in the example subject spectrum in Figure 1C with circles and arrows denoting the difference is power at each end of the filter band. It is difficult to assess the cause of the discrepancy between the FFT and frequency sliding in real data, since the ground truth signal properties are unknown. Thus, we analyzed simulated data wherein an oscillation of 10 Hz with varying amplitude was embedded within a 1/*f^x^* signal with a varying exponent (see Figures 1A and 2). Exponent and amplitude values were chosen to reflect ranges observed in real data (see Methods).

A first analysis explored the impact of varying the aperiodic exponent while keeping the oscillation amplitude (and frequency) constant. Figure 3A shows the results of frequency sliding when there is hardly an oscillation present in the data (amplitude = 0.1), which can occur if a researcher applies the method blindly to a pre-defined band without first inspecting the power spectrum to determine the presence of an oscillation. This analysis shows that, despite the unchanging frequency in the simulated signal, frequency sliding (across time; Figure 3A, left, and averaged over time; Figure 3A, right) underestimates the true frequency (10 Hz), and this bias increases as the aperiodic exponent increases (i.e., the slope becomes more negative). When the oscillation amplitude is large, the bias is reduced (Figure 3B and C show medium and large amplitude oscillations, respectively), however only at very large amplitudes (e.g., 15 microvolts) and/or very shallow slopes does the frequency sliding estimate become unbiased.

The dependency on amplitude makes sense when the data are conceptualized as a combination of periodic and aperiodic features: when the amplitude of the periodic component increases, the contribution of the aperiodic slope to the filtered signal becomes less influential. An implication of this, however, is that, for values of the exponent within the range of real data (around the middle of the simulated range), amplitude changes alone can produce differences in the estimated frequency of the oscillation. This can be seen clearly in the matrix presented in Figure 4, which visualizes the joint effects of amplitude and exponent on the frequency estimated via frequency sliding. Moving along columns within the middle rows, for instance, there is a clear change in frequency driven only by amplitude changes. These results confirm that the underestimation of instantaneous frequency via frequency sliding is due, at least in part, to the bias towards low-frequency power that describes the 1/*f*-like structure of electrophysiological data. This can be further confirmed by simulating an unrealistic but informative case where there is a positive spectrum slope (negative exponent), which causes frequency sliding to *overestimate* the true simulated frequency (not shown).

Lastly, we explored how the specific filter band used here (8 - 12 Hz) might impact the results. If the power spectrum slope drives the observed underestimation because of a difference in power between the lower and upper edge of the filter, than widening the band should lead to an even larger underestimation. This is precisely what was observed (Figure 5A) when the bandwidth of the filter was increased by 6 Hz to span 5 - 15 Hz. This had a dramatic effect (cf. Figure 3) on the underestimation magnitude, which now reached as large as 1.5 Hz lower than the true 10 Hz signal and was never fully attenuated as long as 1/*f* activity was present, even with large amplitude oscillations (Figure 5A). In contrast, using a very narrowband filter (9.5 - 10.5 Hz), strongly attenuated the underestimation, except at steeper slopes. This would be expected since the difference in power at 9.5 and 10.5 Hs is small. And though very narrowband filtering (e.g., 9.5-10.5 Hz) might seem like a solution to overcome estimation bias, the use of such a filter, which highly constrains the signal to within a limited band (i.e., 1 Hz) is antithetical to the premise of frequency sliding, which aims to recover meaningful variation in frequency.

A researcher might unknowingly design a filter that does not fully capture the oscillatory activity under investigation if, for example, they had not inspected individual subject power spectra and detected oscillatory peaks. This scenario was simulated by placing the filter band just below (Figure 5C) or just above (Figure 5D) the 10 Hz peak. A filter that misses the peak will lead to frequency sliding estimates that either strongly underestimate (if below) or strongly overestimate (if above) the simulated frequency in manner dependent on the aperiodic exponent and the oscillation amplitude, with larger exponents and smaller amplitudes generally leader to greater misestimations. Thus, probably more so that slope or amplitude changes in the actual signal, filter bandwidth and position can have a large impact on the estimate obtained via frequency sliding.

Our final set of analyses was aimed at removing the bias by attenuating the aperiodic component of the time series. Although there are several solutions for fitting and removing the 1/*f* feature of a power spectrum, existing methods assume that the subsequent analyses remain in the frequency domain, which means that the power values can be negative. We therefore designed an algorithm to demodulate the 1/*f* shape with minimal impact on the local spectral structure in a way that preserves the amplitude nonnegativity and phase spectrum, which in turn allows for a reconstruction of the signal in the time domain that has minimal 1/*f* (see Figure 6). This demodulation was done separately per trial to allow a diversity of 1/*f* shapes.

We then computed frequency sliding on the demodulated time-domain signals. As in Figure 4, we averaged frequency estimates over time for each combination of exponent and amplitude in the simulated signals. The resulting matrix is shown in Figure 6B on a similar color scale as Figure 4 to allow for visual comparison. The resulting estimates were closer to the ground truth frequency of 10 Hz: the root-mean squared error from 10 Hz on single-trials across all conditions was 0.40 (range across conditions: 0.36 – 0.60) as compared to 0.65 (range: 0.41 – 0.94) for the non-corrected signals. Although the algorithm produced a slight over-estimation of the true frequency (indicated by a max value across the whole matrix in Figure 6B of 10.047 Hz), this bias was an order of magnitude smaller than the maximum underestimation in the non-corrected signals and, most importantly, the bias did not appreciably co-vary with 1/*f* exponent or oscillation amplitude (Figure 6B). Thus, our algorithm serves as a proof-of-principle for reconstructing a time-domain signal with suppressed 1/*f*, which can then be used to obtain less biased frequency sliding results.

## Discussion

Simulations were conducted to understand the circumstances and possible root causes underlying the misestimation of instantaneous frequency by the frequency sliding method (Cohen, 2014). Frequency sliding has been adopted in the field to examine the temporal dynamics in the frequency content of oscillatory brain activity (Gulbinaite, Viegen, et al., 2017; Nelli et al., 2017; Noguchi & Kubo, 2020; Samaha & Postle, 2015; Shen et al., 2019; Wutz et al., 2018), yet the behavior of this method has not been thoroughly investigated in signals where ground-truth properties are known and where physiologically-relevant parameters of the signals are manipulated (e.g., oscillation amplitude and spectral slope). The simulations demonstrate that frequency sliding systematically underestimates the true oscillation frequency when there is 1/*f* structure in the data – a feature of nearly all neural field potentials. In fact, when oscillatory activity (defined as a bump in the spectrum above and beyond the 1/*f*) is absent or very weak, variation in frequency sliding may be due solely to spectral exponent changes. Variation in the slope (exponent) of the 1/*f* activity can readily lead to changes in the recovered frequency, even when the true frequency is constant (Figures 3 and 4). Because spectral slope is known to vary within (Donoghue et al., 2020; Podvalny et al., 2015) and between-individuals (Donoghue et al., 2020; Schaworonkow & Voytek, 2020; Voytek et al., 2015), slope variation could confound the interpretation of estimates obtained via frequency sliding. Moreover, our simulations showed that as oscillation amplitude decreases, the influence of the 1/*f* background on frequency sliding becomes stronger, thus, amplitude changes alone could produce changes in the instantaneous frequency estimated via frequency sliding (Figure 4), presenting another possible confound.

The choice of filter bandwidth and location also has an impact on the estimated frequency (Figure 5). The influence of aperiodic features in the data becomes exaggerated as the edges of the filter enlarge beyond the bandwidth of the oscillatory peak. This suggests that using a wide-band filter as a means of being agnostic about the specific frequency of an effect is not good practice. A filter that misses an oscillatory peak, perhaps because a researcher uses a pre-defined filter band for all subjects or because they are simply unaware of the location of a peak, also leads to systematic amplitude- and exponent-dependent misestimations of the true oscillation frequency. This result highlights the complex interaction between oscillation amplitude, aperiodic exponent, and filter choices that should be considered when interpreting frequency sliding results.

The observation that 1/*f* activity will skew the energy present in the filtered signal is a general principle that has implications for spectral analyses beyond frequency sliding. For instance, 1/*f* activity may also influence any frequency estimation method that relies on the distribution of spectral energy across a band, such as the center-of-gravity method (Klimesch et al., 1993), though this would need to be systematically explored. Moreover, when extracting broadband activity from a signal, such as when investigating broadband gamma activity (e.g., 70 - 150 Hz), the use of a single wide-band filter spanning the whole range of interest will predominantly reflect only the lower frequencies and should be avoided. Instead, multiple separate estimates of smaller sub-bands within the range should be obtained (via multiple filters or wavelet convolution, for instance) and individually normalized prior to being combined.

### Assumptions and limitations

The simulation parameters used here were meant to mimic aspects of real data. This necessitates certain assumptions that should be considered in order to fully contextualize the results. A fundamental assumption is that the physiological processes that underlie oscillation amplitude and frequency modulation are independent of those that generate the aperiodic activity. Although there is not currently clear evidence one way or the other, this should be noted since a violation of this assumption could imply that empirical cases should not be expected to match the simulated conditions. A close relationship between the 1/*f* and the narrowband dynamics would also mean that the anti-1/*f* demodulation process might remove meaningful signal.

Moreover, real power spectra often contain multiple spectral peaks and features beyond the 1/*f* component and oscillatory parameters. For instance, spectra can be additionally parameterized with an offset (addition of energy at all frequencies) component and a ‘knee’ (a region of flatter slope typically at slow and infra-slow frequencies (Donoghue et al., 2020). Exploration of how variation in these features may contribute to frequency sliding would be interesting, though the general principles observed here are not expected to differ. However, one surely important feature of spectral slope changes is the ‘rotation’ frequency about which the slope varies. This was not explored here, but one might expect that if a frequency band under consideration were further or closer from the rotation point, the impact of slope and/or amplitude on frequency estimates may vary. This would suggest caution when comparing frequency sliding results across different brain areas, which have different rotation points (Podvalny et al., 2015), or across different frequency bands, which may be differently positioned with respect to the rotation point. On the other hand, electrode- and trial-specific 1/*f* demodulation might help mitigate this effect. Further simulations would be required to determine these complex relationships.

If a filter range is centered on the oscillatory peak and has a bandwidth that matches that of the oscillation, then slope and amplitude interact to impact frequency estimations by, at most, 0.2 Hz (Figured 3 and 4). One may question whether 0.2 Hz is a large enough bias to worry about. The answer depends, in part, on the effect size (in Hz) that one deems theoretically meaningful. In most examples in the literature where frequency sliding has been applied to non-invasive data, significant differences in frequency sliding have been an order of magnitude *smaller* than 0.2 Hz. For instance, trial-to-trial changes in alpha frequency of, on average, 0.04 Hz were found to significantly predict accuracy in a two-flash visual discrimination task (Samaha & Postle, 2015) and a difference of ~0.05 Hz predicted accuracy in an orientation discrimination task (Nelli et al., 2017). Even modulating the luminance of a stimulus, which has a large impact on sensory responses, produced a maximum difference of around 0.06 Hz in the alpha band (Cohen, 2014). Only in electrocorticography recordings, which presumably suffer less from signal mixing, significant alpha frequency differences of between 0.5 and 1 Hz were found to predict perceptual reports in a bi-stable motion stimulus (Shen et al., 2019). Note that EEG recordings from the same paradigm replicated the effect but with a difference of about 0.06 Hz (Shen et al., 2019). As our results show, however, if the filter bandwidth used to estimate frequency sliding is modestly broader than the oscillation peak, the misestimation could be as much as 1-2 Hz (Figure 5). Even with a more optimal filter, changes in oscillation amplitude or 1/*f* slope could be sufficient to produce frequency effects of a comparable magnitude to those reported in the literature. This means that researchers using the method should take several steps to help rule out the influence of oscillation amplitude and aperiodic slope.

### Recommendations

First, it can be helpful to assess whether the temporal dynamics of frequency modulation are really important for the hypothesis under consideration. If peak frequencies can be compared between conditions using spectrum-based methods (peak finding, gaussian fitting, etc.), this would side-step issues associated with frequency sliding. If temporal dynamics are important, then the following recommendations might be useful when using frequency sliding.

#### 1. Always plot spectra

At a minimum, researchers should inspect power spectra across conditions of interest. If aperiodic spectral features differ between conditions or if oscillatory amplitude differs between conditions, this could be cause for cautiously interpreting frequency differences. Plotting spectra in log-log space can help visualize slope changes. Even better, spectral properties can be explicitly parameterized (Donoghue et al., 2020) and statistically compared. If amplitude or slope differences are found, the values reported from the present simulations may serve as a rough guide to assess whether those differences are sufficiently large to cast doubt on the frequency effect.

#### 2. Check spectral peaks

If there is no peak in the power spectrum, there is likely no oscillation, and frequency sliding results will be driven by spectral slope or data features other than frequency shifts. Visualizing spectral peaks can also provide a secondary analysis to validate any frequency sliding differences (as done in Wutz et al., 2018, for example). Although keep in mind that the frequency precision of an FFT of a short data period can be limited.

#### 3. Carefully design the filter

A filter with a plateau-shaped frequency response is key for mitigating bias toward the center of the filter band (Cohen, 2014). In addition, the simulations here suggest that centering the filter on the oscillation peak and ensuring that the bandwidth matches closely the oscillation bandwidth can help protect against biases. This should be done on subject-by-subject level (and on a frequency-by-frequency level, if different bands are being studied). A recently developed method to determine empirical frequency-band boundaries (Cohen, 2021) may further help to define appropriate numerical boundaries for the filter.

#### 4. Attenuate the 1/f activity

We presented a method to attenuate the 1/f component in the frequency domain, and then reconstruct the signal in the time domain (to which frequency sliding can then be applied). This approach may not perfectly remove the bias if the polynomial fit does not capture all of the 1/f shape, however, it will help reduce the bias when a strong aperiodic component is present in the data. Our approach to removing the 1/*f* and reconstructing the time-domain signal could, in principle, be combined with more sophisticated algorithms for estimating the shape of the aperiodic component of the power spectrum (e.g., the FOOOF algorithm (Donoghue et al., 2020)) so long as the anti-1/*f*-modulator was normalized or constrained to produce non-zero values in the demodulated amplitude spectrum. This approach may provide even better results than in our simulations, which used a relatively simple polynomial fitting routine.

#### 5. Apply a spatial filter

There are several spatial filtering methods that will construct a component out of multichannel data based on maximizing the energy in a narrow frequency range, relative to the broadband energy (Cohen, 2017; Haufe et al., 2014). Such an analysis may further help isolate narrowband from broadband activity, thereby reducing the slope of the 1/f.

### Summary

Frequency sliding is a useful tool to extract time-varying frequency information from neural signals, but users should know how the spectral properties of their data interact with frequency estimation. Through simulation, the present work highlights the potential for oscillatory amplitude and aperiodic slope changes to confound frequency sliding estimates. This, along with an exploration of how filter properties can impact the results of frequency sliding analyses, should motivate researchers to better understand underlying sources of estimated frequency variation in their data.

